# Close encounters of the three morphs: does color affect aggression in a polymorphic lizard?

**DOI:** 10.1101/2020.05.02.074146

**Authors:** Stefano Scali, Marco Mangiacotti, Roberto Sacchi, Alan Jioele Coladonato, Mattia Falaschi, Luca Saviano, Marina Giulia Rampoldi, Matteo Crozi, Cesare Perotti, Francesco Zucca, Elisabetta Gozzo, Marco Alberto Luca Zuffi

## Abstract

Color polymorphism is genetically controlled and the process generating and maintaining morphs can affect speciation and/or extinction rates. Competition and aggression among morphs can contribute to polymorphism maintenance and color badges are useful signals in intraspecific communication, because they convey information about alternative strategies and avoid unnecessary conflicts. This could lead to an uneven spatial distribution of morphs in a population, because the local frequency of each morph establishes the intensity of the competition in that neighborhood, and then the fitness of each male. We used a polymorphic lizard, *Podarcis muralis*, to assess if aggression varies among morphs under two contrasting hypotheses: a heteromorphic vs. a homomorphic aggression. We used laboratory mirror tests after lizard color manipulation and we verified the results consistency with the analysis of the spatial distribution of morphs in a wild population. Both the experiments confirmed that aggression is morph-specific and notably homomorphic. The adoption of behavioral alternative strategies that minimize risks and costs of unwanted conflicts could facilitate the stable coexistence of the phenotypes and reduce the resource competition. A bias in aggression to like-coloured males would advantage rarer morph, which would suffer less harassment by common morphs and obtain a fitness advantage. This process would be negatively-frequency-dependent and would stabilize polymorphism in the populations, possibly leading to sympatric speciation.

## Introduction

Color polymorphism occurs when two or more heritable color morphs “*coexist in temporary or permanent balance within a single interbreeding population […] in such frequencies that the rarer cannot be due solely to mutation*” (Huxley 1955). Morphs are genetically controlled and can evolve by both natural and sexual selection, and the process generating and maintaining it can affect speciation rates and/or extinction rates, either positively or negatively (Gray and McKinnon 2007; Hugall and Stuart-Fox 2012). In several species of polymorphic lizards, morphs generally associate with alternative reproductive strategies, modulated by complex interactions among environmental pressures (e.g. social interactions and individual density), and therefore represent locally adapted optima (Sinervo and Lively 1996; Svensson et al. 2001; Roulin 2004; Sacchi et al. 2007a). When males of different morphs associate with opposite breeding strategies, the intensity of the aggressive behavior may sensibly diverge among morphs, and color may reliably predict the outcome of a dyadic encounter, irrespective of other asymmetries in size, residency or prior experience (Hover 1985; Thompson and Moore 1991a; Thompson and Moore 1991b; Sinervo and Lively 1996). For example, green males of the tree lizard (*Urosaurus ornatus*) are more likely to dominate orange ones despite their lower size (Hover 1985); orange-throated males of side-blotched lizards (*Uta stansburiana*) are highly aggressive and dominant over the other morphs (Sinervo and Lively 1996), as well as red males of the Australian painted dragon (*Ctenophorus pictus*) are more likely to win dyadic contests with yellow males (Healey et al. 2007). However, in other lizard species, the role of color morph in intra-sexual contests is quite controversial, if not completely ineffective to predict the outcome of combat between males of different morphs (Stuart-Fox and Johnston 2005; Sacchi et al. 2009). For example, color morph does not predict the contest outcome or aggression levels for two species of dragon lizards, even if a marginal effect may subsist only during fight between unfamiliar opponents (Stuart-Fox and Johnston 2005).

One of the mechanisms that can help to maintain variation in a population is negative frequency-dependent selection resulting from some processes, such as predation (Olendorf et al. 2006; Qvarnström et al. 2012), sexual selection (Sinervo and Lively 1996) and intraspecific competition (Seehausen and Schluter 2004), that guarantee an advantage to rare morphs. Morphs often adopt alternative strategies that are an outcome of male-male competition and maintain genetic variation and enhance the reproductive success of each morph under context-dependent control (Bleay et al. 2007; Hurtado-Gonzales and Uy 2010). In this scenario, males of rare morphs should receive less aggression from other males, because they do not share the same resources, and thereby they should experience a fitness advantage (Seehausen and Schluter 2004; Qvarnström et al. 2012). The spatial scale at which behavioral interactions among morphs confers a fitness benefit is the composition of morph within a neighborhood, because the local frequency of each morph establishes the intensity of the competition in that neighborhood, and consequently the fitness of each male (Zamudio and Sinervo 2003). When morphs associate to alternative breeding strategies, the fitness of a given male depends on the number of males that come in direct competition with him, and consequently on the relative frequency of morphs in its neighborhood. For example, the three male morphs of the side-blotched lizard (*Uta stansburiana*) exhibit a combination of alternative breeding strategies that interact in a rock-paper-scissor game (Sinervo and Lively 1996). Orange males are aggressive and defend a territory, blue-males are aggressive, but are mate-guarder, and yellow males are not aggressive and behave like sneakers. Each morph has specific behavioral traits that allow it to outcompete only one of the other two. So, orange males outcompete yellow males but are defeated by blue males. There is experimental evidence that the frequency-distribution of morphs in a neighborhood of each male directly affects its fitness, in a way that orange males in a neighborhood of blue males achieve much less fitness than orange males in a neighborhood of yellow males (Zamudio and Sinervo 2003). Therefore, the analysis of the spatial distribution of morph at the neighborhood scale can reveal information on the competitive interactions among the strategies behind morphs.

Furthermore, morphs could reduce intraspecific competition by divergence in resource use (i.e., a character displacement) and this process is acknowledged as one of the possible mechanisms for sympatric speciation (Qvarnström et al. 2012; Svensson 2017) and makes the study of color polymorphism particularly attractive for evolutionary researchers.

The Common wall lizard (*Podarcis muralis*) is a small (50–70 mm adult snout–vent length, SVL) diurnal lizard of central and south-eastern Europe (Sillero et al. 2014), whose males vigorously fight for territories, showing a marked territorial behavior (Edsman 1990; Sacchi et al. 2009) and which express a pigment-based ventral color polymorphism in both sexes with three discrete color morphs (white, yellow, and red, Sacchi et al. (2007b); Sacchi et al. (2013)). Morphs are genetically determined (Andrade et al. 2019) and correlations between morphs and aggressiveness remain controversial (Sacchi et al. 2009; Abalos et al. 2016; Coladonato et al. 2019). In a previous study using a resident-intruder design (Sacchi et al. 2009), we showed that simple rules such as residency and body size differences determine the outcome of agonistic encounters but we did not find any effect of color on male aggression or fighting success. By analyzing dyadic encounters in a neutral arena, Abalos et al. (2016) found that the black patches emerged as a good predictor of contest outcome independently of morphs, even if red males lost fights against heteromorphic males more often than yellow or white males. However, this effect could be due to a correlation with the size of black patches. The plasmatic concentration of testosterone also did not differ among morphs (Sacchi et al. 2017), but only on the basis of the season as yellow males maintained significantly higher T-levels over time and displayed a stronger subsequent decline. The hormone profile did not differ between red and white males (Sacchi et al. 2017). Accordingly to seasonal variation in hormone profile, Coladonato et al. (2019) were able to detect morph-specific differences in the seasonal pattern of variation in the aggressive behavior of yellow males with respect males of the other two morphs, but not between white and red males. Overall, no clear picture emerges from these studies. Further investigations are, therefore, necessary to clarify the relationship between color morph and aggression in this species, also with a view to a better understanding of the strategies adopted by each morph.

A recent study highlighted the effectiveness of mirror experiments for measuring the intensity of the aggressive behavior in lizards, demonstrating that they do not show self-recognition and attack the mirror image as a true “rival” (Scali et al. 2019). Mirror tests have the great advantage of allowing the experimenters to control for the effect of asymmetries in size and residence/motivation, as well as color signal, since each individual can be acclimated in one cage until it becomes resident before facing an intruder having the same size and giving a positive feedback during aggressive contests (Scherer et al. 2016). In this study, we used mirrors to assess if aggression varies among morph after removing the main determinants of contest outcome in this species (i.e. size and residency, Sacchi et al. (2009)). In detail, we explored two contrasting hypotheses: i) a heteromorphic aggression hypothesis (3MH), where each morph compete with others in a balanced scenario; ii) a homomorphic aggression hypothesis (HH), where each morph shows a higher aggression level when faced with an opponent displaying the same color (i.e. the same strategy). These hypotheses were tested in two ways: i) a laboratory experiment where we used mirrors to test aggression after color manipulation, and ii) an intensive field sampling where we assessed if the spatial distribution at neighborhood level of different morphs supports the results of the laboratory experiment.

## Methods

### Laboratory methods

Thirty-six adult male lizards (12 for each pure morph) were captured by noosing during the reproductive period (April-June 2018) in five sites in the Milan province (northern Italy). Animals were carried to the Natural History Museum of Milan and housed in individual plexiglass boxes (40 × 40 × 30 cm) with a refuge positioned near one box?s wall, water *ad libitum* and fed with three *Tenebrio molitor* larvae/day. A sheet of absorbent paper was used as substrate, so that the terrarium could keep the resident odor and the lizards could consider it as their territory. The vertical sides of the boxes were covered externally with white paper sheets, to avoid external stimuli and wall reflectance.

After three days lizards were tested in the same terrarium where they were acclimated, removing water and food. The ventral color was manipulated painting throat and belly with water-based tempera colors, randomly assigning all the color combinations with four replicas for each combination. The same individual was tested for four consecutive days, maintaining the same coloration used in the first treatment, to verify response consistency. A total of 144 trials were performed during the experiment.

A heating lamp (ZooMed Repti Basking spot lamp, 150 W) was turned on for 15 minutes before each trial to achieve an optimal body temperature similar for all the individuals (Sannolo et al. 2014). A photographic set was then placed on the terrarium providing uniform led lightning and a webcam (Microsoft Life Cam HD-3000) was used to record the trial. The lizard was put under the refuge for one minute while the mirror (30 × 15 cm) was positioned at the opposite side of the terrarium. When the camera was turned on, the refuge was removed and the behaviors were recorded for 15 minutes. All the lizards were released, healthy, at the capture sites after the experiments.

The videos were analyzed using the BORIS (Behavioral Observation Research Interactive Software), an open-source software (www.boris.unito.it (Friard and Gamba 2016)). We counted the number of bites against the mirrored image to quantify the aggressive response to the stimulus. Since some lizards never bit the mirrored image, to prevent zero-inflation problems, the number of bites was transformed to the probability of bites during the trial (BP = no. of interactions with bites/no. of total interactions). We defined “interactions” any time the lizard entered the half of the terrarium bearing the mirror. Individuals who never bit the mirror in any trial were discarded from the following analyses.

### Field methods

Field sampling was performed in the archaeological site of Castelseprio (Province of Varese; UTM 32 T 489077E, 5063874 N; 338 m a.s.l.). The site is on the top of a hill and is characterized by stone ruins in an open area with grass or bare soil surrounded by natural deciduous woods. The study area is 2380 m^2^ wide, and ruins cover about 10% of the whole surface and their height ranges from 30 cm to 5 m. Common wall lizards are scattered on the ruins and along the wood borders of the whole study area. Field sampling was performed by ten researchers on 3^rd^ April 2017, (09:00-18:00), during lizards? reproductive period, intensively searching across the whole area. We restricted the sampling to a single day in order to have a stationary picture of the distribution of lizards in the study area. We captured a total of 255 lizards by noosing, including 206 adults lizards (SVL>56 mm). Lizards were marked with a non-toxic color to avoid pseudoreplication and released after recording their sex, morph, and position. To be sure that the position errors were small enough, compared to inter-lizards distances (e.g., less than 10 cm), lizards were located on a high-resolution orthoimage obtained by the use of a Remotely Piloted Aircraft System (Fig. 1; Supplementary material): the map had mean horizontal and vertical errors of 3.5 and 2.2 cm, respectively. Analyses were performed on 144 adult males, excluding females, juveniles, and three males belonging to the yellow-red morph.

**Figure 1.**
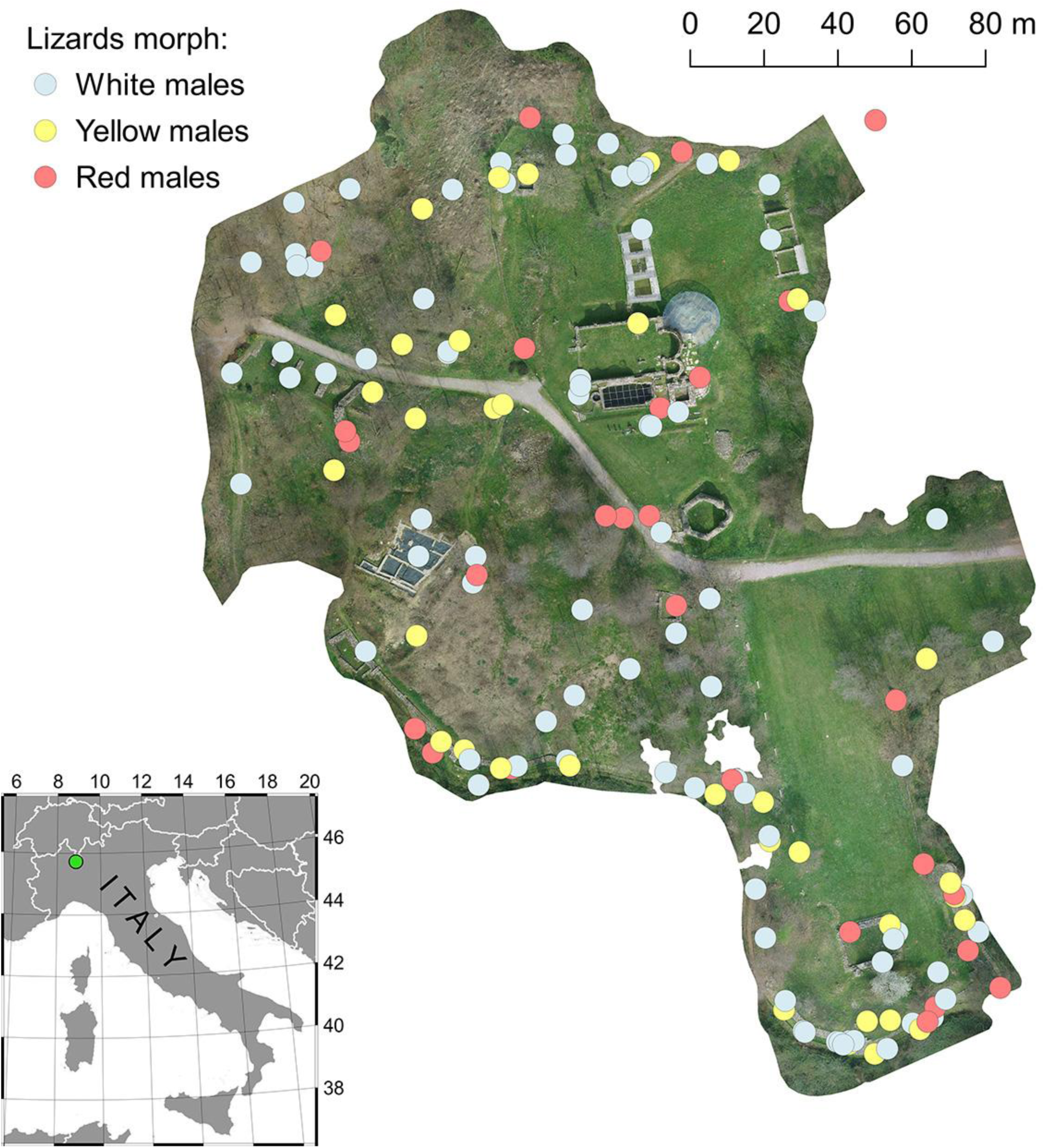
Position of the 144 adult males considered in the analyses. Orthoimage was obtained by the use of a Remotely Piloted Aircraft System, as described in the supplementary material (Menegoni et al. 2019).

## Laboratory statistical analyses

Laboratory data were analyzed using a generalized linear mixed (GLMM) model with a binomial error distribution, using bite probability (BP) as the response variable. For the 3MH hypothesis, we used a model with morph (three-levels factor: W, Y, R), treatment (three-levels factor: W, Y, R), and their interaction as fixed factors, trial (entering as a continuous variable given the constant time between subsequent trials) as covariate controlling for a potential habituation effect (Peeke 1984); the lizard ID was used as a random factor on the intercept to account for repeated trials. For the HH hypothesis, we used a model with treatment (recoded into a two-level factor: homomorphic, heteromorphic) as the fixed factor, trial as the control covariate, and ID as the random intercept.

Predictor significance was evaluated by likelihood-ratio (LR) tests (Zuur et al. 2009). Analyses were performed under the R 3.45.2 statistical environment (R Development Core Team 2018), using the package “glmmTMB” (Brooks et al. 2017).

### Field statistical methods

The same hypotheses (3MH and HH) used for lab experiment were tested on field data. First, we mapped the distribution of all males (Fig. 1) and computed for each focal lizard the distance to: the nearest white (d_W_), the nearest yellow (d_Y_), the nearest red (d_R_), the nearest homomorphic (d_Ho_), and the nearest heteromorphic (d_He_) conspecific. Secondly, we computed the differences in the minimum distance for each morph pair (Δ_RW_ = d_R_ - d_W_; Δ_YW_ = d_Y_ - d_W_; Δ_RY_ = d_R_ - d_Y_) associated to each focal lizard, and between the homomorphic and heteromorphic minimum distances (Δ_HH_ = d_Ho_ - d_He_): this way, we obtained a measure of the relative proximity of males of the three morphs, or of homomorphic and heteromorphic males to each focal lizard. In the end, we averaged these Δs according to the hypothesis to be tested: under 3MH, for each focal color morph we derived the three average, 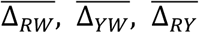; under HH, the average 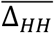 We then tested whether the observed average 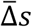 significantly departed from zero, which would have been interpreted as a morph-specific spatial response (repulsion or attraction). Since the number of individuals of the three morphs was not balanced (white = 81, yellow = 36, red = 27), differences in 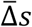 may simply reflect the relative abundance of a given morph (i.e. a rarer morph can show a higher distance just by chance). This is true also for 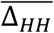even in a scenario where morphs relative abundance were balanced (1/3 white, 1/3 yellow, 1/3 red), the probability of being closer to a heteromorphic would be higher just because of chance. To account for this spatial bias, we adopted a data permutation procedure, and built the expected average 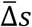 null distribution given the observed point pattern and assuming color morph not to affect the spatial distribution of male lizards. We first permuted lizard color morph (no. of permutations = 999), and we re-computed all the distances (d) and distance differences (Δ). We thus obtained the null distributions of all the average 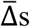 for each focal color morph (3MH hypothesis), and the null distribution of 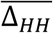 (HH hypothesis). We then assessed the probability of each observed 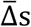 being larger or smaller than the respective null distribution (depending on the observed value being larger or smaller than the mean null value).

## Results

### Laboratory experiment

Out of the 36 lizards tested, six males never responded to the stimulus, i.e., they never moved after refuge removal or they never enter the half of the terrarium bearing the mirror. From the remnant 30 focal lizards we obtained 119 usable trials, one was excluded because the focal lizard did not enter the half with the mirror. So, 29 lizards had four replicas, and one had three.

The 3MH model for BP did not reveal any significant main effect (morph: LR, χ^2^ = 0.359, d.f. = 2; P = 0.836; treatment: LR, χ^2^ = 1.539, d.f. = 2; P = 0.463; morph × treatment: LR, χ^2^ = 5.295, d.f. = 4, P = 0.258; trial: χ^2^ = 3.650, d.f. = 2; P = 0.056). Only “trial” showed an almost significant P-value, with a weak negative effect on BP (β_estimate_ = -0.170; β_SE_ = 0.089). On the opposite, the random effect of the individual identity was highly significant (LR, χ^2^ = 75.018, d.f. = 5; P < 0.001), and accounted for 18.34% of the total BP variance, highlighting the occurrence of a strong variation among individual propensity to bite the mirrored image.

Considering the HH model, trial was confirmed almost significant (LR, χ^2^ = 3.656, d.f. = 1; P = 0.056) and with a similar negative effect (β_estimate_ = -0.170; β_SE_ = 0.089) as 3MH model. The re-coded two-level treatment showed a significant effect, instead (LR, χ^2^ = 5.584, d.f. = 1, P = 0.0181), which highlighted a higher aggression level during homomorphic contests compared to heteromorphic ones (Fig. 2). Again, the random factor was highly significant (LR, χ^2^ = 43.367, d.f. = 1; P < 0.001) and explained 19.80% of the total variance.

**Figure 2.**
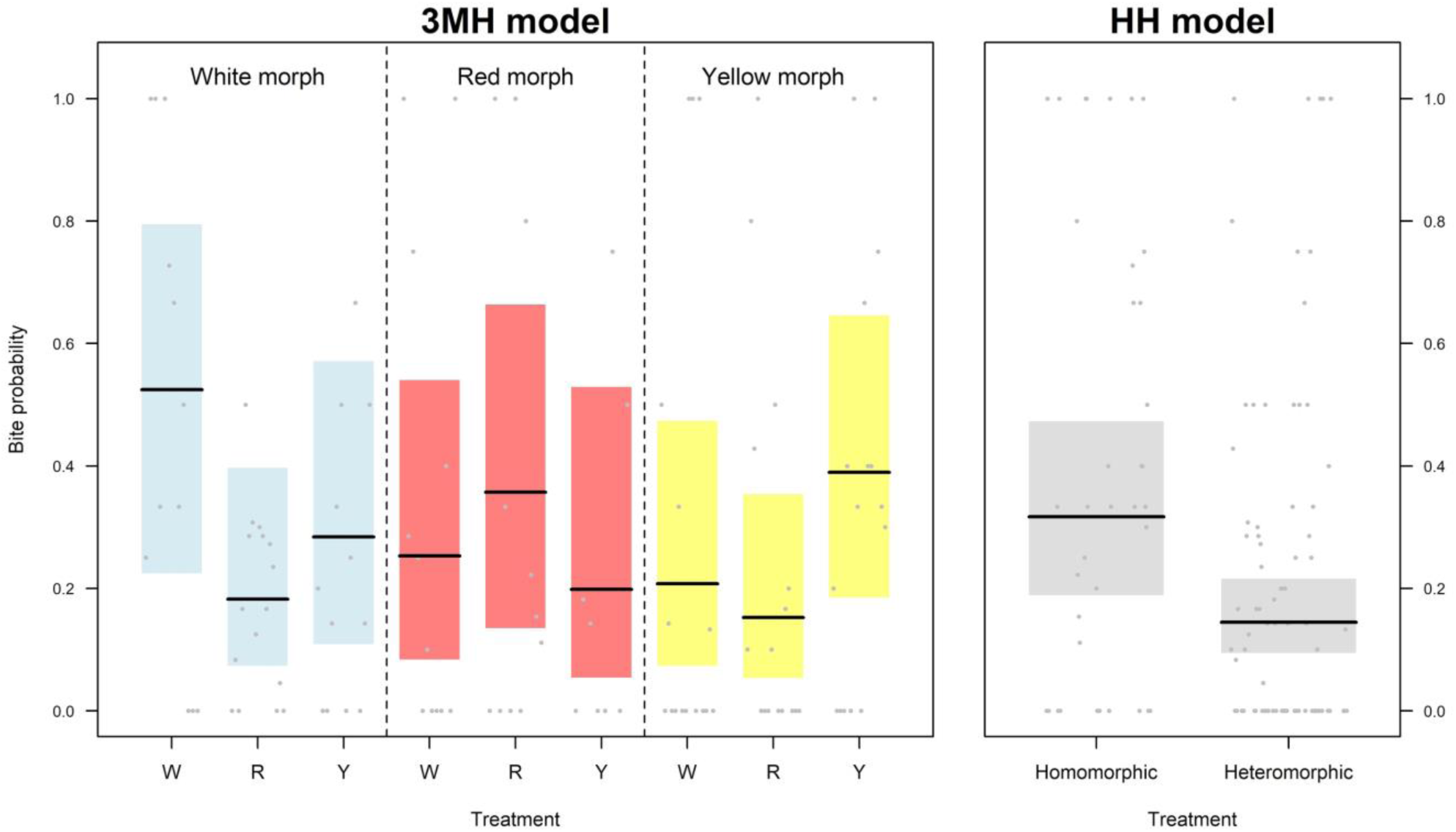
Effect of treatment and morph on the biting probability (BP), according to the heteromorphic aggression model (3MH, left panel), and on the homomorphic aggression model (HH, right panel). For 3MH model, all morph × treatment combinations were shown. Grey points = observed values; horizontal thick lines = predicted BP values; colour or grey shaded area = 95% confidence interval of predictions.

### Field experiment

No difference was observed in the mean distance between individuals when morph pairs were compared (Fig. 3), i.e.: mean distance from red morph is similar to that from white (Δ_RW_ = 7.53 m; P = 0.272) and yellow morph (Δ_RY_ = 1.63 m; P = 0.246); the mean distance from a yellow morph is equal to the distance from a white morph (Δ_YW_ = 5.90 m; P = 0.378). On the opposite, the mean distance between homomorphic and heteromorphic individuals resulted significantly higher than null mean distances (Δ_HH_ = 4.53 m; P = 0.043; Fig. 3)

**Figure 3.**
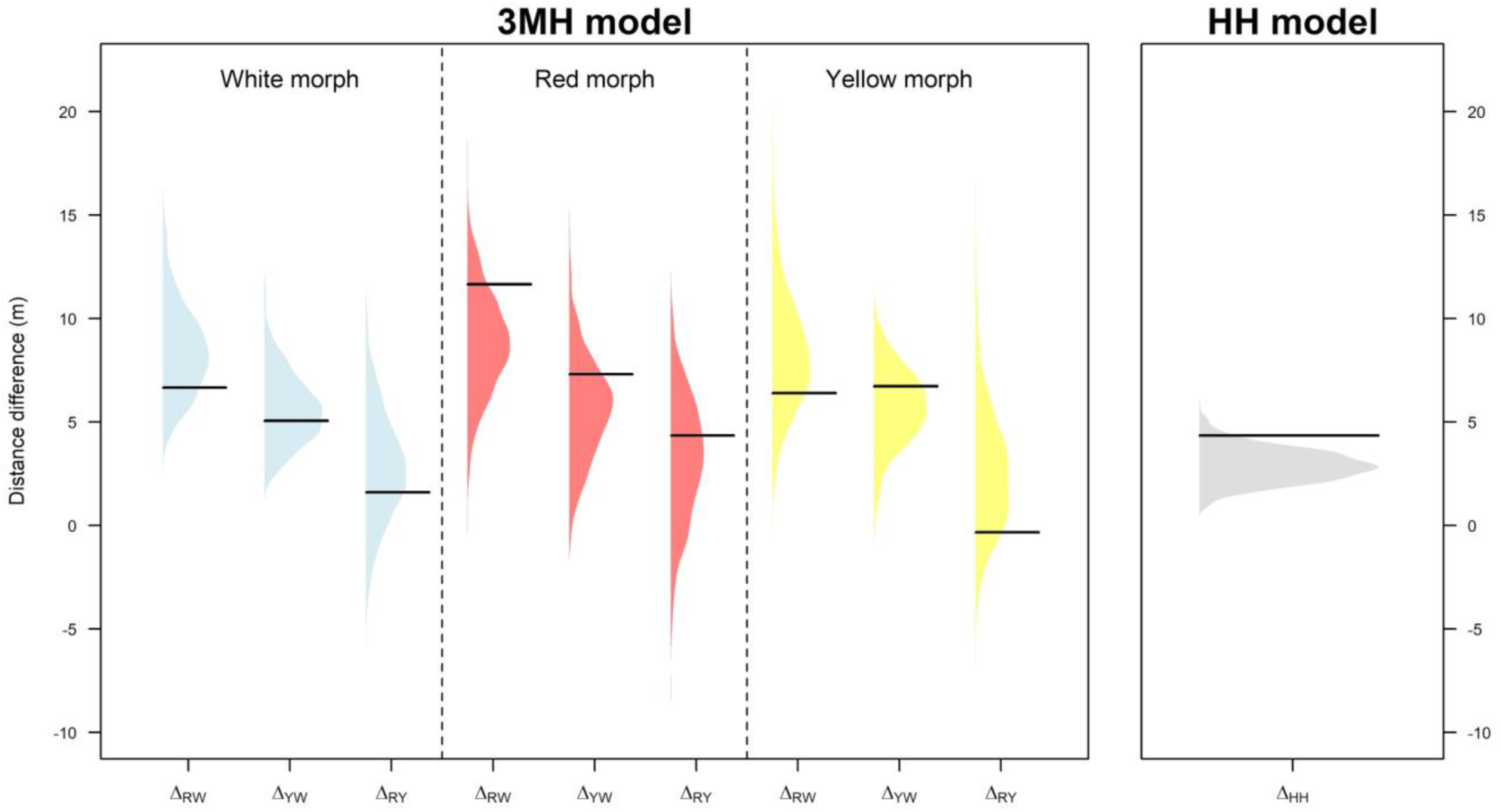
Observed distance differences (black lines) compared to the distribution of the simulated distance differences (color or grey shaded areas). The left panel shows all morph combinations under the 3MH hypothesis, while the right panel shows the average differences between homomorphic and heteromorphic individuals (HH hypothesis).

## Discussion

Our experiment demonstrated that the common wall lizard shows a morph-specific aggressiveness. The balanced experiment based on the three-morphs hypothesis showed that lizards are often aggressive versus intruders of each morph, but no difference was observed in preferential aggression towards a specific morph. On the contrary, even if most of the lizards showed aggressive behaviors towards mirrored opponents, higher aggression was observed towards individuals belonging to their same morph.

Morph-specific aggression has been studied in the past years to understand mechanisms underlying polymorphism maintenance with contrasting results. Many studies were conducted on fish, mainly on cichlids, that are famous for their spectacular intraspecific diversity, and in some cases males compete more heavily with other males of the same color, as in our case study (Seehausen and Schluter 2004; van Doorn et al. 2004; Dijkstra et al. 2007; Dijkstra et al. 2008; Dijkstra et al. 2010). In these cases rare male phenotypes would receive less aggression than common male phenotypes and this could generate frequency-dependent selection, as rare morphs are more likely to gain higher dominance status as a result of reduced harassment from competing males (Dijkstra et al. 2010). Although this mechanism does not contribute to the emergence of reproductive barriers, it can pave the way for sympatric speciation (Dijkstra et al. 2008). Furthermore, if males bias their aggression towards similar rivals that share the same resources, rare morphs should receive less aggression from other males and thereby experience advantage in establishing breeding territories (Seehausen and Schluter 2004; Dijkstra et al. 2007; Qvarnström et al. 2012).

The previous study demonstrated that common wall lizards can discriminate colors and can, consequently, recognize individuals belonging to their own morph (Pérez i de Lanuza et al. 2018). This ability presumably contributes to different results in staged contests, with red males having the lower fighting ability (Abalos et al. 2016), and to assortative mating, with homomorphic male-female pairs more common than the heteromorphic pairs (Pérez i de Lanuza et al. 2012).

Assortative mating could be advantageous also in terms of fitness, since non-random mating produces clutches with different characteristics, in accordance with the r, K and mixed strategies demonstrated for the common wall lizard (Galeotti et al. 2013). Under these assumptions, throat and belly coloration can be considered as a visual badge that common wall lizards use in intraspecific communication. If males preferentially choose homomorphic females, then homomorphic males should be considered as the direct competitors for this resource, so the higher aggression level observed in our experiment can be explained as a need to defend the territories by intruders in a resource competition (Seehausen and Schluter 2004). This conclusion is supported by the lack of differences in aggression in the 3MH model. Similar results were obtained for *Ctenophorus decresii*, an agamid with four discrete morphs (Teasdale et al. 2013), that showed a higher aggressive behavior during homomorphic contests when models were presented to free individuals (Yewers et al. 2016). The color perception of the opponent seems fundamental in triggering aggression also in other species, such as *Urosaurus ornatus*, as demonstrated by experiments where colors were manipulated (Hover 1985).

Our field experiment supported these results, as the mean distance among individuals resulted significantly different when homomorphic and heteromorphic males were compared, in accordance with the HH model. On the contrary, no difference was observed in the 3MH model, highlighting that there is no variation in the mean distance between pairs belonging to the three morphs. This result suggests a non-random distribution of males in the study area, in a sort of “repulsive” effect of homomorphic pairs. A similar result was observed in haplochromine cichlids of Lake Victoria, where breeding territories of individuals of different colors are closer than those of individuals of the same color (Seehausen and Schluter 2004; Dijkstra et al. 2006). The uneven spatial distribution of male common wall lizards could be reinforced by the individual recognition given by scent marks, that contribute to give information about neighbors. In fact, in lacertids femoral secretions convey information about neighbor characteristics, such as size, weight and familiarity (Carazo et al. 2008) and proteins, in particular, give information about the identity and morph in *P. muralis* (Mangiacotti et al. 2019a; Mangiacotti et al. 2019b). Furthermore, lacertid lizards are able to remember the spatial location of scent marks (Carazo et al. 2008), so males could build a spatial map of neighboring rivals. In this way, they can decide which neighbors could exert a major threat to their territories and address aggression against their direct competitors, minimizing both the energetic costs of territory defense and the risks of suffering injuries or predation, according to the paradigm of the “dear-enemy effect” (Ydenberg 1988; Whiting 1999; Carazo et al. 2008; Tumulty 2018).

In conclusion, male aggression in the common wall lizard seems to be morph-dependent. The adoption of behavioral alternative strategies that minimize risks and costs of unwanted conflicts could facilitate the stable coexistence of the phenotypes (Dijkstra et al. 2006; Yewers et al. 2016). A bias in aggression to like-colored males would advantage rarer morph, which would suffer less harassment by common morphs and obtain a fitness advantage. This process would be negatively-frequency-dependent and would stabilize polymorphism in the populations (Seehausen and Schluter 2004; Dijkstra et al. 2006; Dijkstra et al. 2008).

## Acknowledgments

This research was carried out in conformity with current European and Italian laws on animal use in scientific research (authorized by the Ministero dell?Università e della Ricerca, MIUR, prot. 0002154/PNM, 2016, 2nd March, valid over 2016-2018) and Italian laws for the access to the Archeological Sites (authorized by the Ministero dei Beni e delle Attività Culturali e del Turismo, prot. n. 1441/MIBACT-SAR-LOM, 2017, 17th March We are grateful to all the students who helped us during the field work, in particular: Carlotta Pasquariello, Mara Battaiola, Cristian Matellini, Simone Buratti, Francesca Baccalini, Manuela Springolo, Sara Pozzi, Thomas Zabbia, Pietro Rodigari, Stefano Pezzi, Ilenia Bresciani,. We also thank Dr. Sara Matilde Masseroli of the Sovraintendenza Archeologica, Belle Arti e Paesaggio per le Province di Como, Lecco, Monza e Brianza, Pavia, Sondrio e Varese for the technical support within the archeological site.

