## Supplementary Materials for "Close encounters of the three morphs: does color affect aggression in a polymorphic lizard?"

### SUPPLEMENTARY MATERIAL

The georeferenced orthoimage was realized by the use of a Remotely Piloted Aircraft System (RPAS) to acquire a series of high-resolution images that were then converted into a 3D model using structure from motion (SfM) software. The RPAS-based digital photogrammetric survey was conducted with the acquisition of 288 digital photographs. The collected images had a minimum overlap and sidelap of about 85% and 80%, respectively. The flights was planned with 50 m AGL (Above Ground Level). The average GSD (Ground Sample Distance) of images is 1,22 mm/px. The survey area is 4.63 Ha.

The features of RPAS platform and on-board camera are reported in Table 1.

Tab. 1. RPAS and on-board camera specifications

| RPAS system specifications | | | | | |
|---|---|---|---|---|---|
| **RPAS type** | **Dimension** | **Engines** | **Rotor Diameter** | **Empty weight** | **Payload** |
| exacopter | 80 x 80 x 35 cm | 6 brushless | 15" (38.1 cm) | 6.9 kg | 2.5 kg |
| **On-board camera specifications** | | | | | |
| **Camera** | **Sensor type** | **Sensor size** | **Image size** | **Pixel size** | **Focal length** |
| Sony alfa 6000 | CMOS | 23.5 x 15.6 mm | 6000 × 4000 px | 4 x 4 µm | 16 mm |

The RPAS was equipped with a GPS and all the points of the Digital Outcrop Model (DOM) were georeferenced in a WGS84 metric coordinate system. Moreover, to obtain a high accuracy model 20 points in the area were measured with a Topcon Hiper-Pro GNSS (13 were used as Ground Control Points – GCPs – and 7 as Check Points – CKPs).

The 3D digital model (DOM) was developed with the Structure from Motion (SfM) technique using Photoscan Professional v.1.2.5 software (Agisoft, 2016), which is widely employed in earth sciences studies (e.g. Goncalves and Henriques, 2015; Casella et al., 2016; Menegoni et al., 2019).

The procedures used during the processing were schematically the following:

*Image pre-processing.* All the images were georeferenced using the coordinates registered by the on-board GPS and the images with blur effects were discarded.

*Image matching, bundle block adjustment and reconstruction of sparse and dense PCs.* Images were aligned using the highest accuracy (full resolution matching) using the pair pre-selection method that takes in account the image positions registered by GPS, then the bundle block adjustment were computed, and the sparse and dense PCs were reconstructed (over 100 million points).

*Mesh creation.* The 3D mesh was constructed setting the surface type as arbitrary, the dense cloud as data source and a face count and selecting the highest number of face count suggested by the software (~ 20 million faces).

*Texture mapping and orthophoto mosaic generation.* The Generic texture mapping mode was used in order to obtain a proper texture for the mesh, developing a texture atlas composed by an orthophoto mosaic with a resolution of 1,24 mm/pixel was generated as TIFF file.

The final absolute accuracy of the orthoimage of the study area was calculated by comparing GCPs and check point coordinates measured by the GNSS data with coordinates of the same points in the models. The comparison shows a satisfying absolute accuracy with mean horizontal and vertical errors of 3.5 and 2,2 cm, respectively.
